# Large Language Model Consensus Substantially Improves the Cell Type Annotation Accuracy for scRNA-seq Data

**DOI:** 10.1101/2025.04.10.647852

**Authors:** Chen Yang, Xianyang Zhang, Jun Chen

**Affiliations:** Department of Statistics, Texas A&M University, College Station, Texas, 77843, USA; Division of Computational Biology, Department of Quantitative Health Sciences, Mayo Clinic, Rochester, Minnesota, 55905, USA

**Keywords:** Cell type annotation, Collective intelligence, Large language models, Multi-LLM consensus, scRNA-seq, Uncertainty quantification

## Abstract

Different large language models (LLMs) have the potential to complement one another. We introduce an iterative multi-LLM consensus framework for annotating single-cell RNA sequencing data. This framework outperforms the best state-of-the-art method by nearly 15% in mean accuracy (77.3% vs 61.3%) across 50 diverse datasets from 26 tissues, encompassing over 8 million cells. By leveraging cross-model deliberation, our framework quantifies uncertainty, identifies ambiguous clusters for expert review, provides transparent reasoning chains, and minimizes the effort and expertise needed for cell type annotation in large-scale studies. Additionally, our framework enables users to seamlessly integrate new LLMs.

## 1 Main

Cell type annotation is a critical step in single-cell RNA sequencing (scRNA-seq) data analysis and is key to the translation of vast cellular data into biological insights.^1^ Early annotation approaches require human experts to manually compare highly expressed genes in each cell cluster with canonical cell type marker genes from the literature, a process that is both time-consuming and requires specialized knowledge. As datasets grow to millions of cells across diverse tissues, the manual approach has become increasingly intractable. Researchers have proposed various computational methods in the past decade, spanning supervised learning (e.g., SingleR^2^), unsupervised learning (e.g., scCATCH^3^), and knowledge-based (e.g., CellAssign^4^) methods. See^5^ for a recent review.

As the annotation accuracy is still far from optimal, especially for those under-studied tissue types, new methods are continuously being developed. Among recent advancements, popV^6^ and GPTCelltype^7^ represent two promising efforts to enhance annotation accuracy. popV is based on the idea of ensemble learning, synthesizing the advantages of individual annotators, while GPTCelltype leverages the power of large language models to interpret marker gene expression patterns and provide biologically informed cell type annotations based on prior knowledge embedded in the LLM’s training data.

Current state-of-the-art methods still face significant limitations. Although popV can integrate multiple machine learning algorithms, the approach is solely data-driven and is unable to exploit the extensive prior biological knowledge about cell-type-specific gene expression. The LLM-based approach, GPTCelltype, which relies on the prior biological knowledge summarized by the specific LLM, is subject to individual model biases and lacks sufficient uncertainty quantification. The fundamental challenge -integrating complex and often contradictory evidence from marker genes with cell type ontology relationships and knowledge of developmental lineages – remains unresolved in the LLM framework.

Pioneered by ChatGPT, multiple other LLMs, including Claude, Gemini, Grok, Deepseek, Qwen, and others, have been developed lately. As each LLM represents a comprehensive summary of existing human knowledge, dependent on the model architecture, the learning algorithm, and the training dataset, we speculate that the performance of LLM-based cell typing will vary significantly based on different LLMs. Indeed, when we compared the cell-typing performance between different LLMs based on the GPTCelltype algorithm, we observed significant variations in accuracy across different cell types (Fig. 1a). This leads us to hypothesize that by leveraging the strengths of individual LLMs, cell type annotation accuracy can be improved significantly. To achieve this end, we developed a multi-LLM consensus framework for cell typing, mLLMCelltype, to systematically integrate multiple LLMs to reduce individual model biases and to enable better uncertainty quantification through structured collaborative reasoning.

**Fig. 1:**
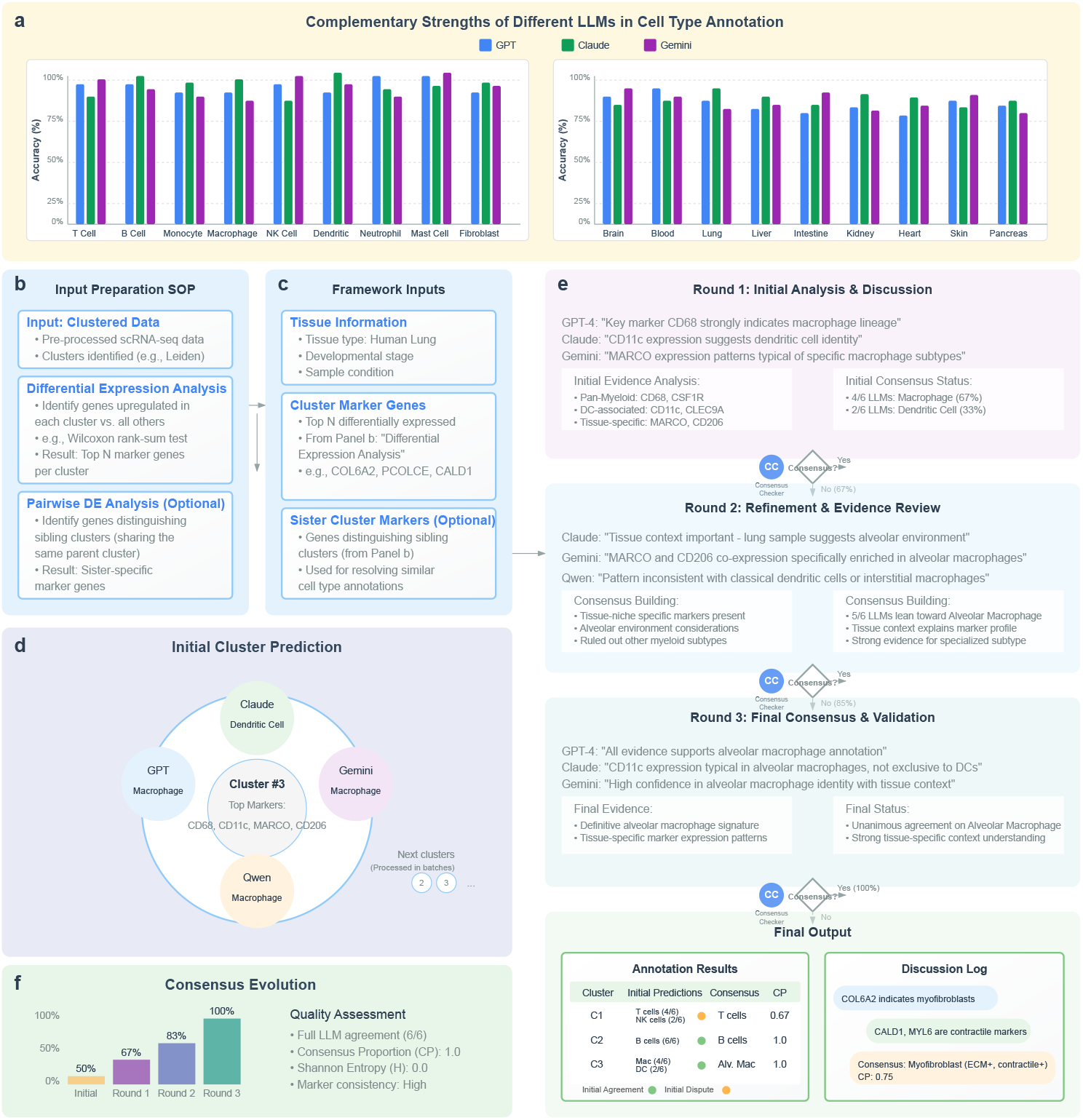
The mLLMCelltype framework workflow. **a**, Complementary strengths: Different LLMs (GPT, Claude, Gemini) show varying performance across cell types and tissues, highlighting the value of a multi-model approach. Performance patterns shown are synthesized from results across the diverse benchmark datasets evaluated in this study. **b**, Input preparation: Differential expression (DE) analysis on clustered data yields marker genes (optional: sister cluster markers). **c**, Framework input: Marker genes (from **b**) and tissue context are provided to the LLMs. **d**, Initial cluster prediction: LLMs independently propose initial annotations per cluster based on the input. **e**, Iterative consensus: LLMs deliberate over multiple rounds, sharing arguments under the guidance of a Consensus Checker (CC) to reach an agreement. Output includes the final annotation and Consensus Proportion (CP). **f**, Consensus evolution: Deliberation progressively improves consensus over successive rounds.

The mLLMCelltype framework integrates multiple LLMs for scRNA-seq cell type annotation, following the workflow illustrated in Figure 1. First, marker genes for each cell cluster are identified through differential expression analysis comparing each cluster against others, derived from pre-processed and clustered scRNA-seq data (Fig. 1b). These marker gene lists, along with tissue context information, form the primary input for our framework (Fig. 1c). Then, multiple LLMs (Fig. 1d) independently receive these inputs and propose initial cell type annotations for each cluster, providing biological reasoning based on the marker gene evidence. For clusters lacking immediate high consensus, the framework initiates an iterative deliberation process (Fig. 1e). During deliberation rounds, the LLMs share their structured arguments, discuss the significance of specific markers (e.g., pan-myeloid CD68 vs. tissue-specific MARCO), consider potential pathway involvement, and evaluate how tissue context influences the cell identity. Each LLM refines its classification by weighing the evidence and reasoning presented by its peers. Crucially, after each round, a dedicated Consensus Checker LLM evaluates the degree of agreement among the participating models against a pre-defined threshold (illustrated by ‘CC’ nodes in Fig. 1e). If consensus is reached, the process terminates, yielding a final annotation and confidence score. If not, deliberation continues for another round (up to a maximum limit), or the cluster is flagged as ambiguous. Figure 1d and 1e illustrate this process using a myeloid cluster example: initial disagreement between ‘Macrophage’ and ‘Dendritic Cell’ based on conflicting markers like CD68 and CD11c (Fig 1d, Fig 1e Round 1) is resolved through deliberation incorporating tissue context (lung) and additional markers (MARCO, CD206), leading to a final consensus of ‘Alveolar Macrophage’ (Fig 1e Round 3). Ultimately, the framework outputs the final consensus annotations, uncertainty metrics (Consensus Proportion -CP), and the reasoning process (Fig. 1e Final Output), with overall consensus typically improving across rounds (Fig. 1f).

Fundamentally, mLLMCelltype achieves enhanced performance through its multi-LLM consensus architecture, which leverages collective intelligence to overcome single-model limitations. Specifically, incorporating a diverse ensemble of complementary LLMs directly addresses the issue of individual model bias; variations in training data and model architecture across LLMs mean that systemic errors or biases present in one model are less likely to be shared by all, leading to a more balanced and accurate consensus (Extended Data Fig. 2). Furthermore, this inherent diversity also fosters knowledge integration, pooling the potentially complementary biological knowledge breadth and depth captured by different models. Crucially, the iterative deliberation process embodies structured reasoning where models critically evaluate differing perspectives. This mechanism actively suppresses inaccurate or unsupported “hallucinations” (Extended Data Fig. 1), a known vulnerability of single LLMs, thereby reducing annotation error rates. The progressive refinement through deliberation leads to a measurable increase in inter-model agreement (Extended Data Fig. 3). Furthermore, the consensus approach inherently improves robustness against noisy input (Extended Data Fig. 4). Together, the bias mitigation through diversity, comprehensive knowledge integration, active error correction via deliberation, and inherent input robustness underpin the superior accuracy and reliability achieved by the mLLMCelltype framework.

We evaluate mLLMCelltype across 50 diverse datasets from different tissues and sources, including Tabula Sapiens (TS), Human Cell Landscape (HCL), Mouse Cell Atlas (MCA), Genotype-Tissue Expression (GTEx) cross-tissue data, Human BioMolecular Atlas Program (HuBMAP) scRNA-seq data, Human Lung Cell Atlas (HLCA), and specialized datasets including B cell lymphoma (BCL) and cancer datasets from colon and lung. We utilize a subset of stable and consistently accessible models: GPT-4o (OpenAI), Claude-3.5-Sonnet and Claude-3.5-Haiku (Anthropic), Gemini-1.5-Pro and Gemini-2.0-Flash-Exp (Google), and Qwen2.5-Max (Alibaba). We focus our comparison to GPTCelltype as GPTCelltype has already been extensively benchmarked against conventional computational methods, demonstrating significantly higher annotation accuracy compared to established tools, including ScType, SingleR, and CellMarker2.0.^7^ The comparison between mLLMCelltype and GPTCelltype highlights the extent to which leveraging multiple LLMs can enhance performance. Across tissue types, mLLMCelltype achieved an average 15% improvement in annotation accuracy (77.3% vs 61.3%) (Fig. 2a) with particularly substantial gains on challenging datasets where GPTCelltype performed poorly. It achieved >95% annotation accuracy on well-characterized cell types and demonstrated significant improvements (20-30%) on datasets that proved challenging for GPTCelltype. For several tissues, our framework demonstrated striking improvements in annotation accuracy: Prostate(GTEx) (86.0% vs 48.0%, +38.0%), Esophagus(GTEx) (87.3% vs 52.5%, +34.8%), and Lung(GTEx) (84.8% vs 52.2%, +32.6%). These tissues present particular challenges due to their complex cellular composition, subtle phenotypic differences between related cell types, and the presence of tissue-specific cell states that require nuanced interpretation of marker gene expression patterns. Notably, in cases requiring deliberation, mLLMCelltype’s ability to track and document the complete reasoning process provided valuable insights into annotation challenges, enabling efficient expert review of contentious cases.

**Fig. 2:**
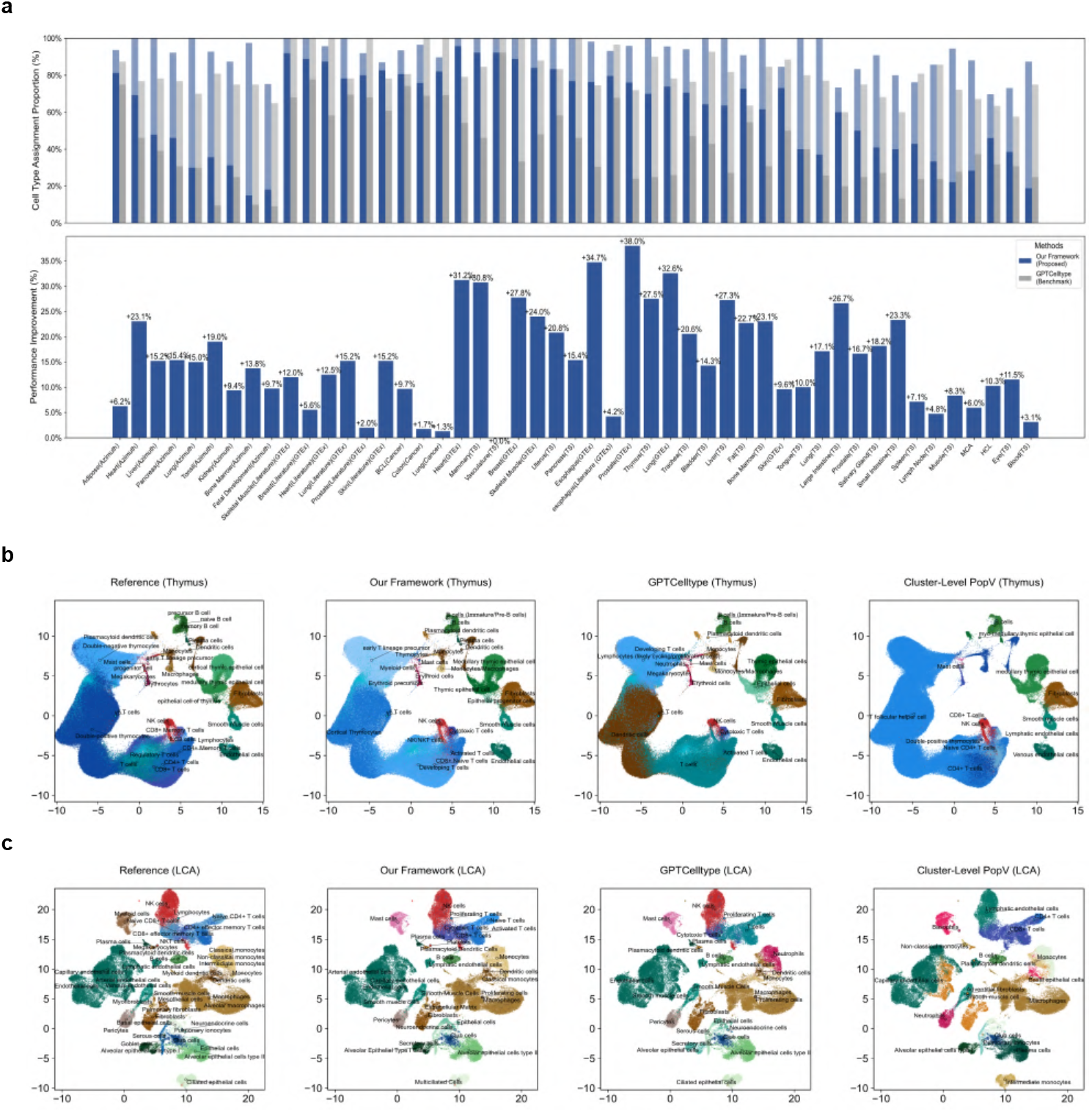
Performance evaluation across diverse datasets. **a**, Comparison with GPTCelltype, showing full and partial matches (see definitions in “Methods”) with accuracy presented as percentages. **b**, Performance comparison on the developmental human thymus cell atlas. UMAP visualizations show cell type annotations from Reference (first), mLLMCelltype prediction (second), GPTCelltype prediction (third), and cluster-level popV prediction (fourth), with points colored by cell type. **c**, Performance comparison on the Lung Cell Atlas (LCA). UMAP visualizations show cell type annotations from Reference (first), mLLMCelltype prediction (second), GPTCelltype prediction (third), and cluster-level popV prediction (fourth), colored by cell type.

We next compare mLLMCelltype to popV, a recent high-performing machine learning-based cell type annotation tool that also employs a consensus-based approach. As popV performs cell-level annotation, cells in the clusters could have different cellular identities. To be comparable, we use the majority rule to achieve cluster-level annotation for popV. We compare the performance on benchmark datasets used in the original popV study.^6^ On the challenging developmental human thymus dataset,^8^ mLLMCelltype demonstrated a 10.6% improvement in annotation accuracy compared to popV (Fig. 2b). On the Lung Cell Atlas (LCA),^9^ mLLMCelltype achieved a 13.17% improvement in annotation accuracy (Fig. 2c), particularly in distinguishing closely related pulmonary cell subsets. Although popV has many unique advantages, such as clustering independence, it faces two major limitations: its reference-based nature and computational demands. popV’s performance heavily depends on the availability of high-quality reference datasets for pre-training, while its matrix-based approach requires substantial computational resources that are not readily available on typical workstations. In contrast, mLLMCelltype requires neither pre-training nor reference datasets, enabling efficient annotation on standard hardware.

While mLLMCelltype does not require reference datasets, it may encounter issues related to training data inclusion. This occurs when the training data includes the benchmark data, which can lead to overestimating the performance. To address this concern, we further validated our method using datasets published after the cutoff dates for LLMs’ training. We evaluated mLLMCelltype on the recently published Human Neural Organoid Cell Atlas (HNOCA),^10^ which integrates 36 single-cell transcriptomic datasets from 26 different organoid protocols, and a comprehensive human peripheral immune cell atlas^11^ that profiles immune cells across 13 age groups spanning the entire human lifespan from birth to over 90 years. Both datasets were published after the training dates of the LLMs used in our framework, providing a rigorous test of generalization to truly new data.

On the HNOCA dataset containing more than 1.7 million cells, our framework achieved high annotation accuracy (96.45% vs. GPTCelltype’s 83.95%; see Extended Data Fig. 5) despite the complexity of neural cell types and developmental states. mLLMCelltype demonstrated particular strength in accurately identifying specific neural progenitor cell types and neuronal subtypes that proved challenging for single-LLM approaches. Similarly, on the lifespan immune cell atlas dataset, comprising 220 healthy donors and extensive multi-omic profiling, mLLMCelltype achieved 78.0% annotation accuracy compared to GPTCelltype’s 65.3% (Extended Data Fig. 6). This dataset is especially challenging due to its fine temporal resolution of immune cell transcriptomes, diverse cell populations, and complex developmental transitions. We successfully validated this capability on even larger datasets, including the HLCA containing over 2.4 million cells (Extended Data Fig. 7).

It is interesting to study the robustness of our framework to noisy marker gene lists. We generated noisy marker gene lists through four types of perturbations: house-keeping gene injection, wrong cell type’s marker gene injection, random gene injection, and random marker gene loss. Notably, mLLMCelltype demonstrates consistent performance even under significant noise, with major cell types maintaining over 90% annotation accuracy up to 30% noise levels across perturbation types (Extended Data Fig. 4). This resilience highlights the framework’s ability to effectively discern biological signals amidst noise. The number of input marker genes is a parameter of our framework. We found that a range between 20-30 genes provides a practical balance between annotation accuracy across different classification complexities and computational cost (Extended Data Fig. 9). Complementing its robustness to input noise, the framework’s cross-model deliberation mechanism also actively mitigates model-specific hallucinations (Extended Data Fig. 1), a feature particularly crucial for accurately annotating rare or ambiguous cell types where single models might speculate. Coupled with the consensus proportion and Shannon entropy metrics derived from the deliberation process, this provides transparent uncertainty quantification, enabling researchers to identify challenging populations requiring expert review – a capability lacking in many automated methods.

Finally, our framework demonstrates consistent performance across different scRNA-seq technologies, showing robust performance on both Drop-seq data^12^ and single-nucleus RNA sequencing (snRNA-seq) data^13^ (Extended Data Fig. 8).

Practically, mLLMCelltype delivers significant advantages for large-scale single-cell analysis. Its modular design allows seamless integration of new LLMs, ensuring future adaptability. The core strength lies in its transparent, systematic consensus process: by tracking the multi-model deliberation, it provides detailed reasoning chains. This empowers experts to pinpoint and review ambiguous cases efficiently based on the consensus metrics, with full context, drastically reducing manual annotation time while enhancing the overall reliability and quality of cell type annotation for complex tissues.

## 2 Methods

### 2.1 Data Pre-Processing and Input Preparation

We processed scRNA-seq data using standard workflows. Raw count matrices were preprocessed using Scanpy^14^ (v1.9.3), including filtering of low-quality cells (minimum 200 genes per cell, maximum 20% mitochondrial reads), normalization (total counts per cell to 1 × 10^4^), log1p transformation, and highly variable gene selection (*n top genes* = 2000). We identified marker genes using differential expression analysis based on Scanpy’s implementation of the Wilcoxon rank-sum test (scanpy.tl.rank genes groups), comparing each cluster against all others. Genes were filtered by log fold change (minimum threshold 0.25) and adjusted p-value (Benjamini-Hochberg correction, *p* < 0.05), then ranked by adjusted p-value to select the top 30 marker genes per cluster. We also performed pairwise comparisons between sister clusters to identify subtype-specific marker genes (see Methods section “Sister cluster analysis”). As an option for our framework, these sister cluster-specific marker genes could improve the annotation resolution (Extended Data Fig. 3).

Our framework accepts marker genes from any differential expression method (e.g., Seurat FindMarkers, DESeq2) and is flexible regarding the number of marker genes provided.

### 2.2 LLM Integration and Deliberation Protocol

We utilized six state-of-the-art LLM instances: GPT-4o (OpenAI), Claude-3.5-Sonnet and Claude-3.5-Haiku (Anthropic), Gemini-1.5-Pro and Gemini-2.0-Flash-Exp (Google), and Qwen2.5-Max (Alibaba). Each model version (Sonnet/Haiku from Anthropic and Pro/Flash from Google) was treated as a separate contributor in the consensus framework. We included different versions from the same providers to increase model diversity and capture complementary strengths in biological knowledge representation and reasoning capabilities. Note that OpenAI’s o1 and DeepSeek-R1 were not included due to limited API availability during our study period. Additionally, models released after our experimental phase (such as Claude-3.7-Sonnet and Grok3) were not incorporated in this analysis. The specific LLM versions used represent their availability at the initiation of our study; future versions or models from other providers can be readily integrated into our framework.

Our framework implements a structured multi-stage consensus-building process: In the initial annotation phase, each LLM receives marker genes for all clusters requiring annotation (processed in batches if necessary) and provides initial cell type annotations based on the marker gene lists. Following this initial phase, the consensus-checker LLM evaluates all clusters to identify those where the consensus proportion threshold has already been met, outputting their annotations directly. For clusters lacking initial consensus, the framework proceeds with an iterative deliberation process, where each LLM is informed of others’ predictions and is asked to provide reasoned arguments for their choices while critically evaluating alternative annotations. This can lead to dynamic opinion shifts as LLMs acknowledge more convincing arguments from their peers. After each deliberation round, the consensus-checker LLM re-evaluates all remaining unannotated clusters to determine if any have reached the consensus threshold, continuing deliberation for those still below the threshold. The number of deliberation rounds is controlled by the `max rounds` parameter, defaulting to three deliberation rounds, empirically determined based on the balance between annotation accuracy and computational cost. This default setting typically achieves satisfactory results while maintaining reasonable API costs. However, users can adjust it to meet their specific needs, increasing the rounds for complex cases requiring extended deliberation or decreasing the rounds when computational resources are limited.

Following each deliberation round, the consensus-checker LLM evaluates whether the consensus proportion threshold has been reached. In our implementation, Claude-3.5-Sonnet serves as the default consensus checker due to its strong reasoning capabilities, with Qwen2.5-Max as a fallback option. Users can customize this selection through the `consensus checker` parameter in our package, allowing them to specify any available LLM as the consensus-checker based on their specific needs. We define “consensus” using a configurable parameter called “consensus proportion threshold” (default: 66.7%, 4/6 in our study), which specifies the minimum proportion of LLMs that must agree on a cell type annotation to be considered a consensus. Once this threshold is reached, the process terminates immediately for the specific cluster, and both the final annotation and the complete reasoning chain are preserved. If the `max rounds` limit is reached without achieving the consensus proportion threshold, the framework outputs the most supported cell type annotation with quantitative uncertainty metrics. Importantly, uncertainty metrics (consensus proportion and Shannon entropy as detailed in Section2.3) are provided for all clusters, regardless of whether they reached the consensus threshold or not. In cases where multiple cell types receive equal support after reaching the maximum number of rounds (a rare occurrence), the framework continues the deliberation process until the tie is broken. If a tie persists despite extended deliberation, the annotation is presented as a composite label (e.g., “celltype1/celltype2”), though such scenarios are extremely rare in practice due to the iterative nature of the deliberation process.

The framework ultimately generates a comprehensive output for each analysis, including (1) final consensus annotations for all clusters, (2) identification of initially controversial versus immediately agreed-upon cases, (3) initial predictions from each LLM, (4) complete deliberation histories for contentious cases, providing transparency into the decision-making process, and (5) quantitative uncertainty metrics for each cluster, including consensus proportion and Shannon entropy values, enabling explicit uncertainty quantification and prioritization of cases requiring expert review.

To ensure structured and effective deliberation, we designed specific prompts for each stage of the process. First, the initial annotation phase prompt emphasizes biological knowledge and marker gene interpretation. Subsequently, the deliberation prompts focus on critical evaluation and biological reasoning, while consensus-checking and summarization prompts are designed to maintain an objective assessment of arguments. The summarization task is executed by the consensus-checker LLM (Claude-3.5-Sonnet by default, with Qwen2.5-Max as an acceptable alternative based on our testing). Detailed prompt templates are provided in Supplementary Table 1.

To ensure broad accessibility and seamless integration with popular single-cell analysis environments such as Seurat and Scanpy, we designed our framework to interface directly with these environments by accepting their specific data objects with marker genes and cluster assignment (e.g., AnnData object for Scanpy, Seurat object for Seurat). Users can create the marker genes using their preferred statistical method (e.g., Wilcoxon rank-sum test, MAST, DESeq2). Our framework processes these marker genes through the LLM consensus pipeline and returns annotation results in compatible formats that can be directly incorporated back into these analysis environments, enabling researchers to continue downstream analyses without disruption to their workflow. This integration approach allows our framework to serve as a complementary component within existing annotation pipelines while significantly improving accuracy and providing uncertainty quantification.

### 2.3 Uncertainty Quantification

Our framework quantifies uncertainty in cell type annotation using two complementary metrics: consensus proportion and Shannon entropy. Both metrics leverage consensus dynamics among multiple LLMs during the deliberation process. These metrics are defined as follows:

#### 1. Consensus Proportion

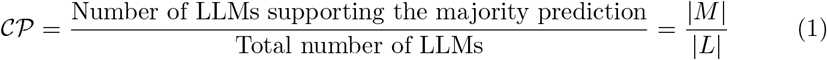

where *M* ⊆ *L* represents the subset of LLMs supporting the majority cell type prediction. With six LLMs, a consensus proportion of 𝒞 𝒫 = 4/6 = 66.7% indicates moderate uncertainty, while 𝒞 𝒫 = 6/6 = 100% reflects complete model consensus.

#### 2. Shannon Entropy

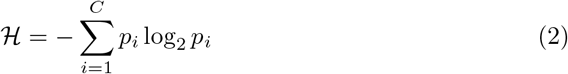

where *C* is the number of unique cell types predicted by the LLMs, and *p*_*i*_ is the proportion of LLMs predicting cell type *i*. Shannon entropy ranges from ℋ = 0 (perfect agreement) to ℋ = log_2_ *C* (maximum disagreement).

These metrics provide complementary information about prediction uncertainty. While Consensus Proportion directly measures the level of consensus, Shannon Entropy quantifies the diversity in opinions among the LLMs. Shannon Entropy is maximized when competing opinions are evenly distributed among LLMs. Clusters with low consensus proportions (default: 𝒞 𝒫 ≤ 66.7%) or high Shannon entropy (default: ℋ > 1) are flagged as ambiguous for expert review, with complete deliberation histories preserved (Fig. 1e).

### 2.4 Hierarchical Annotation Strategy

While our framework’s core functionality focuses on cell type annotation at a single clustering resolution (as described in Figure 1), we also provide an optional extension for hierarchical annotation across multiple granularity levels. This extension is specifically designed for datasets with inherent hierarchical organization, such as comprehensive cell atlases and large-scale reference datasets.

The hierarchical strategy operates by leveraging information from multiple clustering resolutions and specialized marker gene sets. Users need to generate a hierarchical clustering structure, which can be accomplished through:

1. Running Leiden clustering at progressively finer resolutions (our recommended approach) 2. Using established hierarchical clustering algorithms that produce nested clusters 3. Manually defining a biologically meaningful hierarchy based on domain knowledge

For each cluster at each level, two types of marker genes are calculated: 1. Global marker genes: comparing each cluster against all others (standard approach) 2. Sister marker genes: comparing clusters that share the same parent cluster (novel approach)

The framework then applies a sequential annotation process formalized in Algorithm 1. Importantly, this hierarchical extension is provided as a separate workflow from the core mLLMCelltype functionality, requiring additional user input to define the hierarchical structure and generate appropriate marker sets.

#### Algorithm 1

Hierarchical Cell Type Annotation

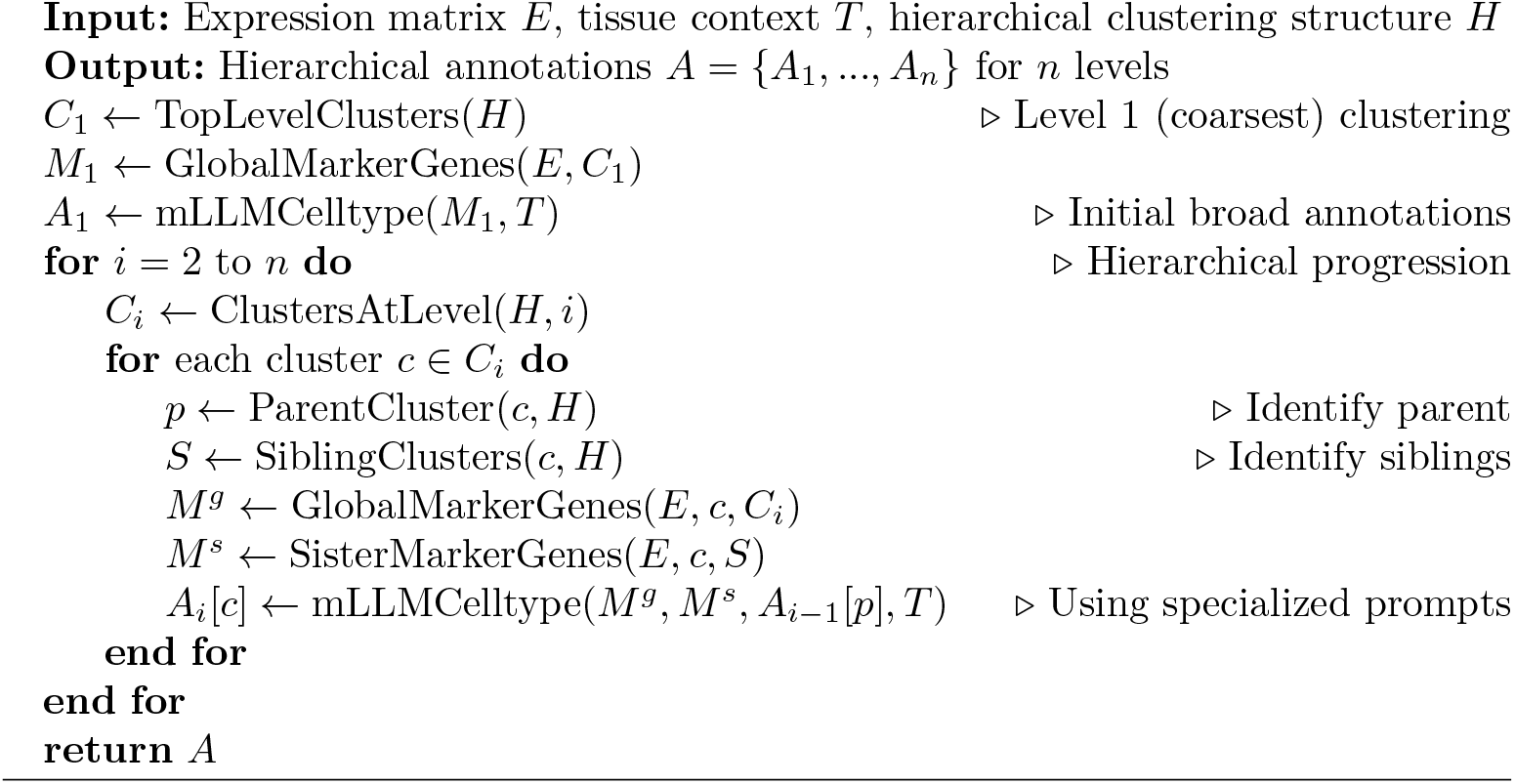

To facilitate this hierarchical approach, we developed specialized prompts that incorporate both global and sister marker information, along with parent cluster annotations (see Supplementary Table 2 for these hierarchical annotation prompts). These prompts explicitly instruct the LLMs to consider the parent cell type as context while evaluating the discriminative power of sister markers.

A key aspect of our hierarchical strategy is how we communicate the hierarchical context to the LLMs through carefully crafted prompts. For example, when annotating the HLCA dataset, we include specific information about the annotation hierarchy and the current level being analyzed:

“Background of Human Lung Cell Atlas (HLCA):

HLCA is a comprehensive integration of 49 datasets of the human respiratory system, combining over 2.4 million cells from 486 individuals. This atlas provides consensus cell type annotations with matching marker genes, including rare and previously undescribed cell types.

Annotation Hierarchy:

Level 1: Basic cell lineages (e.g., immune cells, epithelial cells, endothelial cells, stromal cells)

Level 2: Major cell lineage classifications

Level 3: Specific cell type lineages

Level 4: Defined cell types

Level 5: Cell subtypes

We are currently working on Level 4 annotations -Defined cell types. Please provide cell type annotations at Level 4, focusing on specific cell states and functional subtypes. For example, if you see markers indicating T cells, annotate as ‘naive CD8+ T cells’, ‘memory CD4+ T cells’, or ‘cytotoxic CD8+ T cells’ rather than just broadly as ‘CD8+ T cells’ or ‘CD4+ T cells’.”

This level-specific guidance helps the LLMs understand the expected granularity of annotation at each hierarchical level. Additionally, when analyzing a specific cluster, we provide the parent annotation as context:

“The parent cluster for this group of cells was annotated as ‘T cells’ at Level 3. Now we need to determine the specific T cell subtype at Level 4 based on the provided marker genes.”

We validated this hierarchical strategy on comprehensive atlas datasets, including the HLCA dataset, achieving 89% annotation accuracy at Level 3 annotations with 98% parent-level consistency. The strategy proved particularly effective for rare cell populations, where sister marker genes and parent context helped resolve ambiguous cases. Sister marker genes play a crucial role by providing discriminative features specifically between closely related subtypes. While global marker genes often highlight broadly expressed genes that define major cell types, they frequently fail to capture subtle differences between closely related subtypes. Extended Data Fig. 3 demonstrates that incorporating sister marker genes improved annotation accuracy by 4.8%, particularly in challenging cases where global markers alone were insufficient to distinguish between functionally distinct but transcriptionally similar subtypes, such as different subpopulations of alveolar macrophages or various T cell subtypes within the lung.

### 2.5 Performance Evaluation

To quantitatively assess our framework’s performance, we adopted a unified scoring framework based on Cell Ontology (CL) term relationships:^15^ **Full match** (1.0) for identical terminology and CL terms; **Partial match** (0.5) for shared hierarchical relationships or aligned broad categories; and **Mismatch** (0.0) for annotations with no shared CL ancestry or divergent classifications. The annotation accuracy is calculated as the weighted average of these scores: Accuracy = (*N*_full_+0.5 × *N*_partial_)/*N*_total_, where *N* represents the number of cells in each category. This framework, initially established in GPTCelltype,^7^ provides a standardized way to assess annotation accuracy while accounting for the hierarchical nature of cell type relationships. For a comprehensive comparison with popV,^6^ we converted their predictions to this standardized metric to ensure fair and consistent evaluation across all methods.

### 2.6 Robustness Analysis

To systematically evaluate the framework’s robustness, we designed four types of marker gene perturbations (systematic modifications to marker genes to test framework robustness):

1. **Housekeeping Gene Injection:** Replacing a proportion of cell type-specific marker genes with ubiquitously expressed genes (e.g., *GAPDH, ACTB, B2M*). This perturbation tests the framework’s ability to identify and discount non-discriminative genes, which are often present in statistically derived marker genes due to their high expression levels.
2. **Wrong Cell Type Label Injection:** Introducing marker genes specific to other cell types (e.g., adding B cell marker genes to T cell clusters). This represents the most challenging form of noise as it provides explicitly misleading information, simulating scenarios where statistical methods may identify marker genes shared between related cell types.
3. **Random Gene Injection:** Replacing marker genes with randomly selected genes from the transcriptome, simulating technical noise in marker gene identification that commonly occurs due to batch effects or technical variations.
4. **Marker Gene Random Loss:** Randomly removing a proportion of true marker genes, simulating scenarios with incomplete marker gene information -a common situation in real-world analyses where some genuine marker genes may be missed due to technical limitations or stringent statistical thresholds.

For each marker gene perturbation type, we tested noise levels from 0% to 50% in 10% increments. The noise level represents the proportion of original marker genes affected by the perturbation. Performance was evaluated using our standard scoring framework based on Cell Ontology term relationships. To assess variability, we performed 50 independent trials at each noise level, recording annotation accuracy, deliberation rounds required, and decision types.

### 2.7 Baseline Method Settings

For GPTCelltype comparison, we used their official R package (https://github.com/Winnie09/GPTCelltype, v1.1.1). The differential expression results from our Scanpy pipeline were formatted to match GPTCelltype’s input requirements, and tissue context was provided through the tissuename parameter to ensure the biological relevance of the annotations.

For popV comparison, we used their official implementation (https://github.com/YosefLab/popV) with default parameters. The pre-trained weights were obtained from their HuggingFace model repository (https://huggingface.co/popV). For datasets without available pre-trained weights, we used their recommended “retrain” mode with default settings. For cluster-level popV results, we assigned each cluster the most frequent cell type annotation among individual cells within that cluster, providing a consensus-based representation of popV’s performance at the cluster level. Experiments were conducted using either an NVIDIA A100 80GB GPU or a MacBook Pro with an M2 Max chip (32GB RAM), depending on the computational requirements of specific analyses.

### 2.8 API Cost Analysis

The financial cost of running our multi-LLM consensus framework varies depending on dataset size and the proportion of contentious clusters requiring multi-round deliberation.

GPT-4o: Input: $0.01/1K tokens Output: $0.03/1K tokens

Claude-3.5-Sonnet: Input: $0.003/1K tokens Output: $0.015/1K tokens

Claude-3.5-Haiku: Input: $0.001/1K tokens Output: $0.005/1K tokens

Gemini-1.5-Pro: Input: $0.0005/1K tokens Output: $0.0025/1K tokens

Gemini-2.0-Flash-Exp: Input: $0.001/1K tokens Output: $0.005/1K tokens

Qwen2.5-Max: Input: $0.0016/1K tokens Output: $0.008/1K tokens

Note that these prices reflect rates at the time of our study and are subject to change. LLM pricing typically fluctuates over time, with a general trend toward lower costs following the release of newer models.

For a detailed cost breakdown, we analyzed expense patterns across different dataset scales:

1. **Small datasets** (5-15 clusters): Initial annotation phase averages 2K input/1K output tokens per model per dataset. For datasets where 30% of clusters require deliberation (typical rate), total cost ranges from $0.75-$2.25.
2. **Medium datasets** (16-50 clusters): Processed in 2-3 batches, with deliberation needed for approximately 25% of clusters. The total cost typically ranges from $2.50-$5.00.
3. **Large-scale atlases** (>50 clusters): For datasets like HLCA or Tabula Sapiens, processing requires 4-6 batches with deliberation for 20-25% of clusters. Complete annotation costs between $6.00-$12.00.

Per-cluster costs demonstrate efficiency: direct consensus clusters cost approximately $0.05-$0.08 each, while contentious clusters requiring three deliberation rounds average $0.35-$0.45 each. These estimates include all models’ contributions, summarization, and consensus evaluation.

Cost optimization strategies include (1) using Claude-3.5-Haiku or Gemini-1.5-Pro for the initial annotation phase and summarization stages to leverage their lower token rates, (2) caching deliberation summaries for similar cluster types, and (3) implementing early stopping when consensus emerges before reaching maximum deliberation rounds. These approaches can reduce total costs by 30-40% with minimal impact on annotation accuracy.

While these API costs represent a real expense, they offer significant cost-effectiveness when compared to alternatives. Manual annotation by domain experts typically requires several hours to days of specialized labor per dataset, with costs potentially reaching thousands of dollars for large atlases when accounting for expert time. Similarly, generating new reference datasets for transfer learning approaches requires substantial wet-lab resources and sequencing costs. Our framework provides a favorable cost-benefit ratio by dramatically reducing annotation time while maintaining high accuracy, making it an economically viable solution for both research and clinical applications.

### 2.9 Limitations and Potential Collective Bias

Although our multi-LLM consensus framework significantly reduces single-model hallucination, it may still produce a “collective error” if the models share similar training biases or converge on an erroneous rationale.

Importantly, our current uncertainty quantification metrics (consensus proportion and Shannon entropy) are primarily designed to identify disagreements among models, which is highly effective at flagging ambiguous cases. While our framework significantly reduces annotation errors through multi-LLM consensus, there remains a small theoretical possibility of “collective error” in rare instances where all models might share similar training biases and converge on an incorrect consensus. For example, when analyzing tumor microenvironment single-cell data, a cell cluster expressing both macrophage markers (CD68, CSF1R, CD14, CD163), dendritic cell markers (CD11c, FLT3, ZBTB46), low levels of T cell genes (CD3D, FOXP3), and tumor-specific markers (SPP1, MMP9, VEGFA) might be unanimously but incorrectly classified as “M2 macrophages” with high confidence (CP=100%) by all LLMs. This could occur because all models share similar outdated training data that simplistically categorize CD163+SPP1+ macrophages as “M2-type,” while missing recent research on specialized “mixed-phenotype macrophages” that acquire dendritic and immunoregulatory features in certain tumor microenvironments. However, our extensive benchmarking across diverse datasets demonstrates that such scenarios appear uncommon, with the framework achieving high accuracy across tissue types. Crucially, even for the small minority of potentially challenging cases where collective error might be a concern, the framework’s uncertainty metrics (CP and ℋ) and detailed reasoning chains empower researchers to efficiently target expert review towards the most ambiguous or unexpected annotations. This targeted verification strategy preserves the significant time-saving benefits offered by our automated approach compared to exhaustive manual review, while ensuring optimal accuracy, particularly for novel findings or critical applications.

Future research directions should focus on developing more sophisticated methods to detect potential collective errors. This could include metrics to identify anomalous consensus patterns, such as suspiciously rapid and unanimous agreement on rare or complex cell types that typically require extensive deliberation. For example, if all LLMs immediately agree on a rare subtype without any deliberation, this unusually strong consensus might itself be flagged as a warning sign warranting additional scrutiny. Additionally, integrating structured external knowledge bases (e.g., Cell Ontology, CellMarker database) through retrieval-augmented generation (RAG) or context-augmented generation (CAG) approaches, potentially utilizing a Model Context Protocol (MCP) interface for dynamic database queries, could enable automated cross-validation of LLM consensus against established cell type definitions and marker profiles. These techniques would allow the framework to dynamically access and incorporate specialized biological knowledge during the annotation process.

Another promising direction is developing higher-order reasoning evaluation systems inspired by self-consistency approaches^16^ that analyze patterns across multiple deliberation paths to identify potential errors. Such systems could learn to recognize when LLM deliberations exhibit suspicious patterns that correlate with annotation errors, potentially identifying subtle signals of collective bias even when traditional uncertainty metrics suggest high confidence. These approaches could significantly enhance the framework’s ability to detect and mitigate collective errors while maintaining the efficiency benefits of the consensus-based approach.

## 3 Data Availability

All analyses in this study were performed using publicly accessible datasets.

Comprehensive cellular atlas datasets: The TS dataset was accessed via the UCSC Cell Browser platform (https://cells.ucsc.edu/?ds=tabula-sapiens). The HCL dataset was retrieved from figshare (https://figshare.com/articles/dataset/HCLDGEData/7235471), accompanied by expert annotations from the original study.^17^ The MCA expression matrices were obtained through figshare (https://figshare.com/s/865e694ad06d5857db4b), with corresponding annotations and differential gene lists accessed from the original publication’s supplementary materials.^18^

Tissue-specific atlas datasets: The HLCA data was accessed through the CELLxGENE platform (https://cellxgene.cziscience.com/e/9f222629-9e39-47d0-b83f-e08d610c7479.cxg/).^19^ The LCA dataset was obtained from the CELLxGENE collections platform (https://cellxgene.cziscience.com/collections/5d445965-6f1a-4b68-ba3a-b8f765155d3a). The developmental human thymus cell atlas dataset,^8^ containing over 250,000 cells spanning from childhood to adulthood, was accessed through CELLxGENE (https://cellxgene.cziscience.com/collections/de13e3e2-23b6-40ed-a413-e9e12d7d3910). The HNOCA^10^ was accessed through CELLxGENE (https://cellxgene.cziscience.com/collections/de379e5f-52d0-498c-9801-0f850823c847). The lifespan-wide human peripheral immune cell atlas dataset^11^ was accessed through Synapse (https://www.synapse.org/Synapse:syn61609846).

Cross-tissue and reference datasets: The HuBMAP single-cell data and corresponding annotations were obtained through the Azimuth portal (https://azimuth.hubmapconsortium.org/). For the GTEx analysis, we accessed the gene expression matrices from the GTEx database (https://gtexportal.org/home/datasets), while the expert-curated cell type annotations and differential gene lists were retrieved from the supplementary materials of Eraslan et al.^20^ Literature-derived marker gene sets were also sourced from the same study’s supplementary materials.

Technology-specific and disease-related datasets: The Drop-seq^12^ and snRNA-seq^13^ datasets were obtained by subsetting the HLCA data based on the study field. The BCL dataset was obtained from Zenodo (https://zenodo.org/record/7813151). Cancer-related datasets were accessed through GEO, including colon cancer (accession: GSE132465) and lung cancer (accession: GSE131907) expression matrices and their corresponding cell type annotations. Cell type annotations and marker genes for the non-model mammal analysis were extracted from the supplementary materials of Chen et al.^21^

## 4 Code Availability

The mLLMCelltype package (v.1.0.0) is provided as an open-source software package under the MIT license, with implementations available for both Python and R environments. It is designed to integrate seamlessly with standard single-cell analysis workflows, interfacing directly with Scanpy (via AnnData objects) in Python and Seurat (via Seurat objects) in R. A detailed user manual, example datasets, and step-by-step tutorials are available in the GitHub repository at https://github.com/cafferychen777/mLLMCelltype. This permissive license allows for unrestricted use, modification, and distribution of the software for both academic and commercial purposes. All codes to reproduce the presented analyses are publicly available in the GitHub repository at https://github.com/cafferychen777/mLLMCelltypeReproducibility.

## 5 Extended Data

**Extended Data Figure 1:**
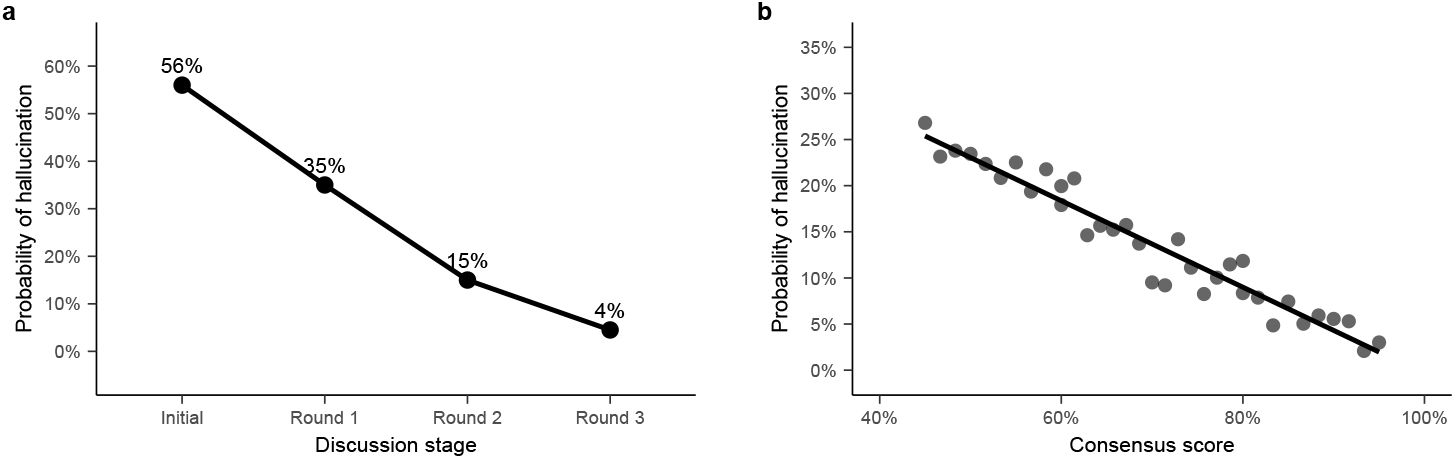
Relationship between model consensus proportion and hallucination reduction during iterative deliberation. **a**, Progressive reduction in hallucination probability across deliberation rounds. In the context of cell type annotation, hallucination refers to when an LLM generates cell type annotations or biological interpretations that are either 1) not supported by the evidence from marker genes, 2) inconsistent with established cell type hierarchies in the literature, or 3) biologically implausible combinations of cellular features. For example, claiming a cell is both a T cell and a B cell simultaneously or assigning neuron-specific marker genes to immune cells would be considered hallucinations. The probability of such events decreases substantially from the initial predictions to the final round of structured deliberation, demonstrating the effectiveness of our iterative consensus-building approach. **b**, Negative correlation between consensus proportion and hallucination probability. A higher consensus proportion among models strongly correlates with lower probabilities of hallucination. This relationship suggests that cases where models achieve high model consensus are particularly reliable, with minimal hallucination risk. Conversely, a low consensus proportion indicates potentially problematic predictions that may require additional deliberation or expert review.

**Extended Data Figure 2:**
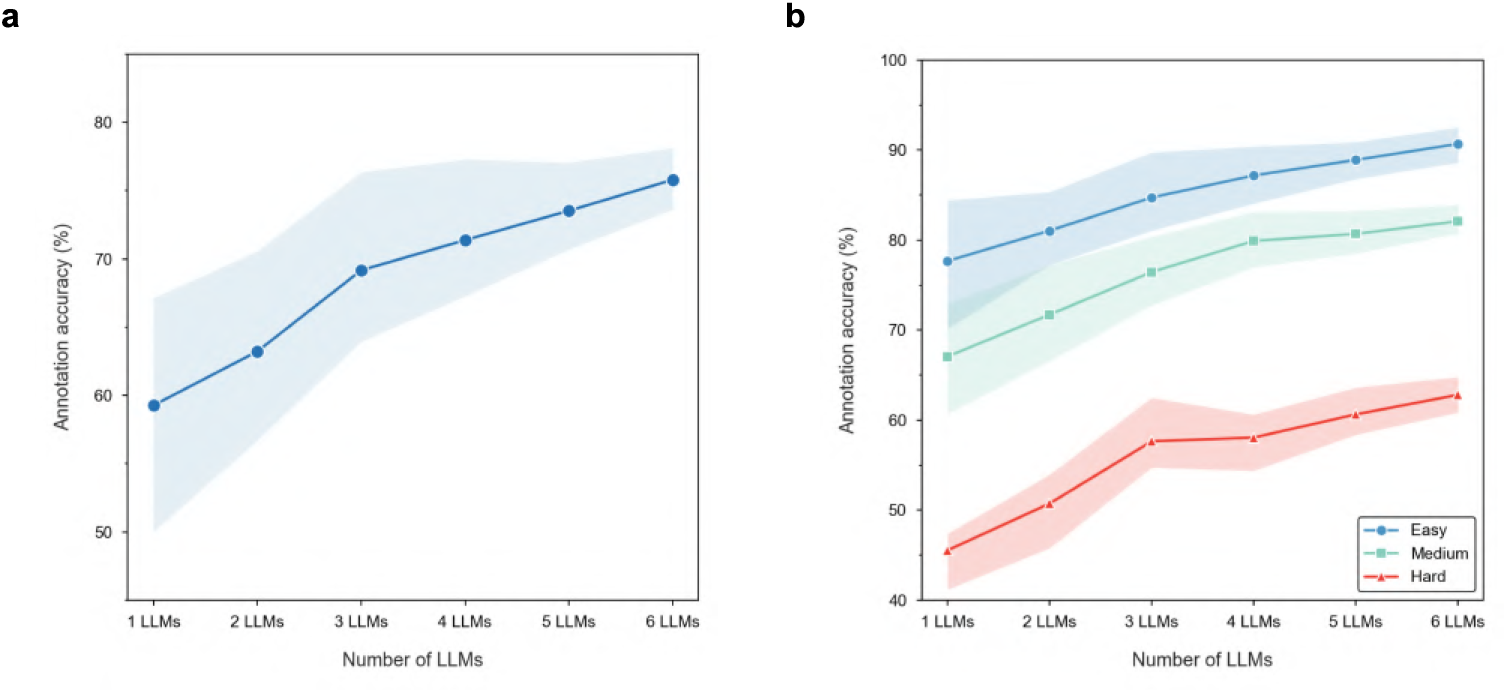
Impact of multiple language models on cell type annotation accuracy. **a**, Relationship between the number of language models and overall annotation accuracy. The line plot demonstrates how cell type annotation accuracy improves as more language models are incorporated into the ensemble, with a consistent upward trend from 1 to 6 LLMs. The shaded area represents the confidence interval based on 100 independent replications, showing reduced variance as more models contribute to the consensus annotation. This demonstrates that ensemble approaches can significantly enhance annotation reliability. **b**, Analysis of annotation performance across different sample difficulty levels. The plots show how annotation accuracy varies for easy (blue), medium (green), and hard (red) samples as the number of LLMs increases. Easy samples achieve over 90% accuracy with 6 LLMs, while medium and hard samples show more substantial relative improvement (22.4% and 38.0% respectively) compared to using a single LLM. This analysis reveals that ensemble approaches are particularly beneficial for challenging cell types that are difficult to annotate with a single model.

**Extended Data Figure 3:**
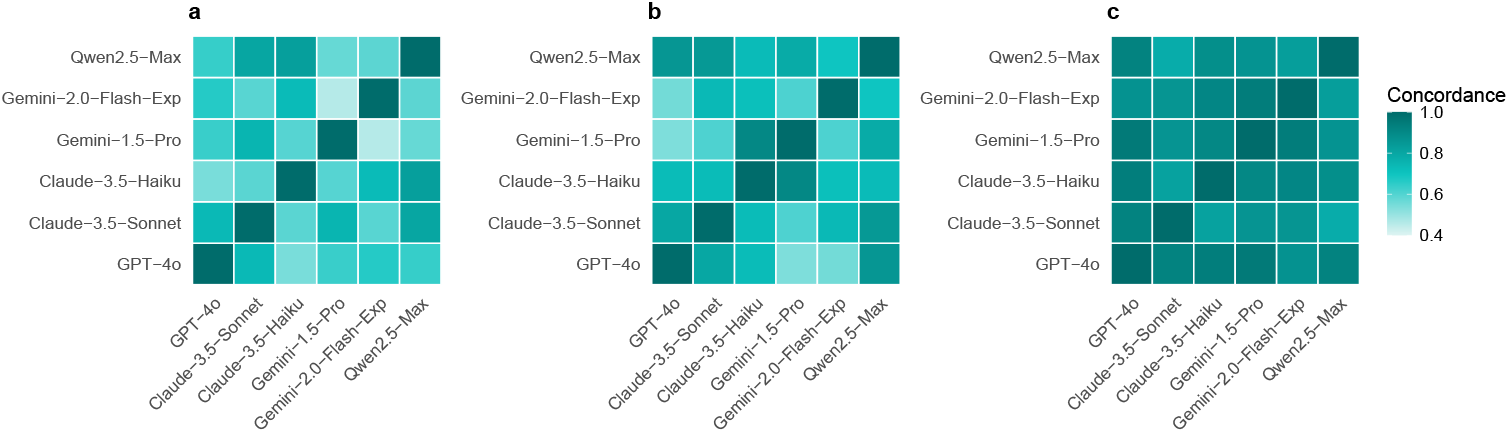
Progressive improvement in LLM prediction consensus proportion across deliberation rounds. **a**, Heatmap showing initial consensus proportion between different LLMs in round 1 predictions. **b**, Heatmap showing increased consensus proportion between LLMs after round 2 deliberation. **c**, Heatmap demonstrating high consensus proportion achieved in round 3, indicating successful consensus building.

**Extended Data Figure 4:**
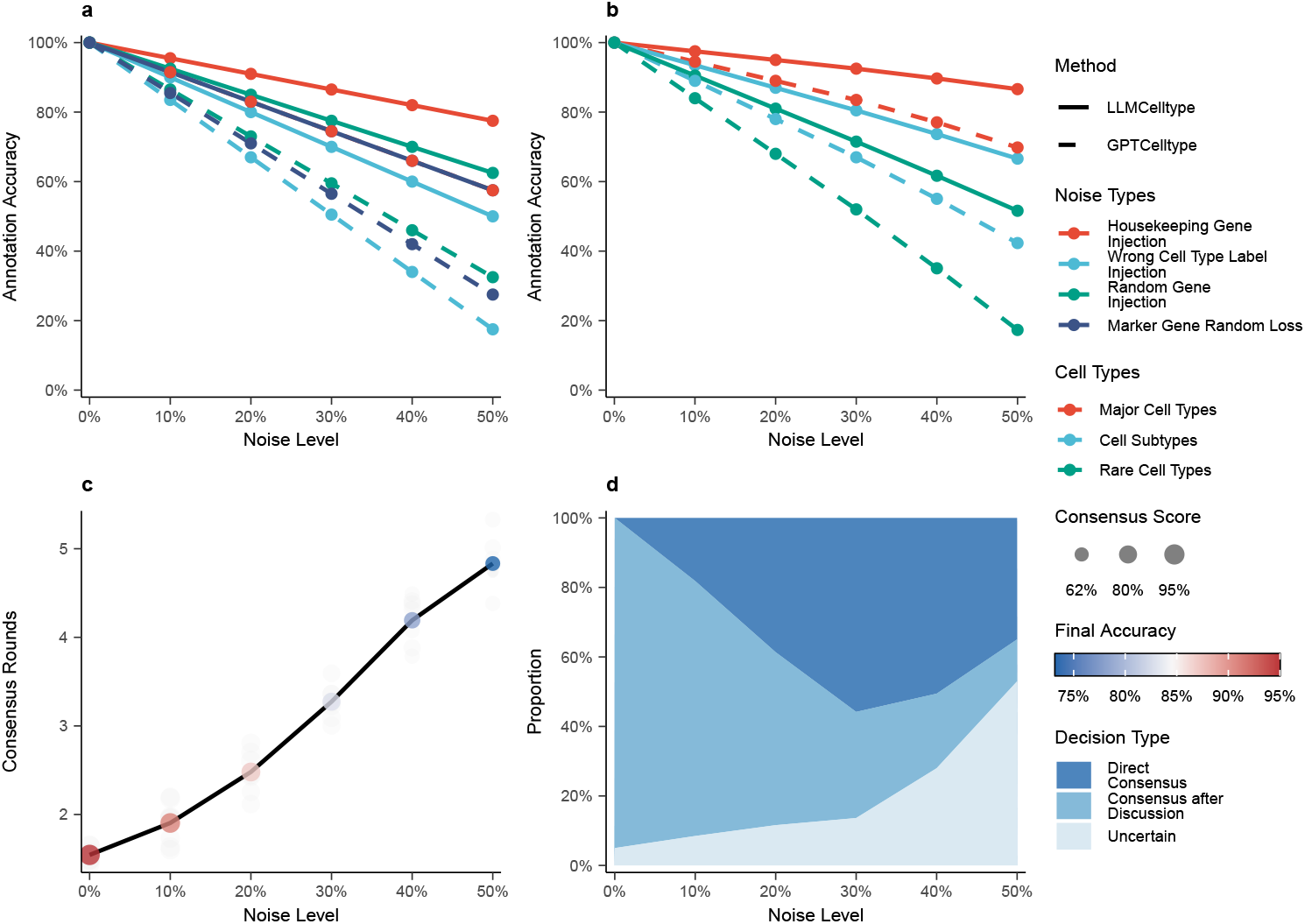
Systematic evaluation of marker genes perturbation robustness in the multi-LLM consensus framework. **a**, Quantitative assessment of annotation robustness under different types of marker genes perturbations comparing our framework and GPTCelltype methods. Our framework exhibits superior robustness patterns across all perturbation types: for housekeeping gene injection, our framework shows only a 4.5% accuracy decline per 10% noise levels compared to GPTCelltype’s 8.5% decline, demonstrating our framework’s enhanced ability to identify non-discriminative genes. For wrong cell type marker injection, our framework (10% decline per 10% noise levels) significantly out-performs GPTCelltype (16.5% decline), indicating our framework’s greater resilience to direct interference with cell type determination. Similar performance advantages are observed for random gene injection (7.5% vs. 13.5% decline) and marker gene loss (8.5% vs. 14.5% decline), validating our framework’s superior ability to prioritize biologically relevant signals over GPTCelltype. **b**, Cell type-dependent robustness analysis reveals distinct performance patterns between our framework and GPTCelltype across cellular hierarchies. For major cell types, our framework maintains >90% accuracy up to 30% noise levels with minimal decay (2.5% per 10% noise levels), while GPTCelltype shows steeper degradation (5.5% decay rate). This performance gap widens for cell subtypes (6.5% vs. 11% decay) and becomes most pronounced for rare populations, where our framework’s 9.5% decay rate substantially outperforms GPTCelltype’s 16% decay rate. This hierarchical response pattern demonstrates our framework’s enhanced capability to leverage redundant marker genes for accurate classification, particularly for challenging rare cell populations where marker gene information is more limited. **c**, Quantitative characterization of LLM c2o1nsensus dynamics under increasing marker genes perturbation. The deliberation requirements scale non-linearly with noise levels, from 1.5 deliberation rounds at baseline to 4.8 deliberation rounds at 50% noise levels, with consensus proportion decreasing from 95% to 62%. Background points represent individual consensus attempts (*n* = 8 per noise level), demonstrating increased variance in deliberation patterns at higher noise levels (standard deviation increasing from 0.1 to 0.35). **d**, Analysis of the framework’s adaptive decision-making behavior using area plots. The distribution of decision types shifts from predominantly direct consensus (95% at baseline, exponential decay rate*−* 0.045) to increased reliance on consensus achieved through deliberation rounds (peaking at 30% noise levels) and flagging uncertainty. Under severe noise conditions (> 30% noise levels), the framework exhibits a marked increase in uncertainty flagging (non-linear growth with exponent 1.8), reflecting its built-in quality control mechanisms.

**Extended Data Figure 5:**
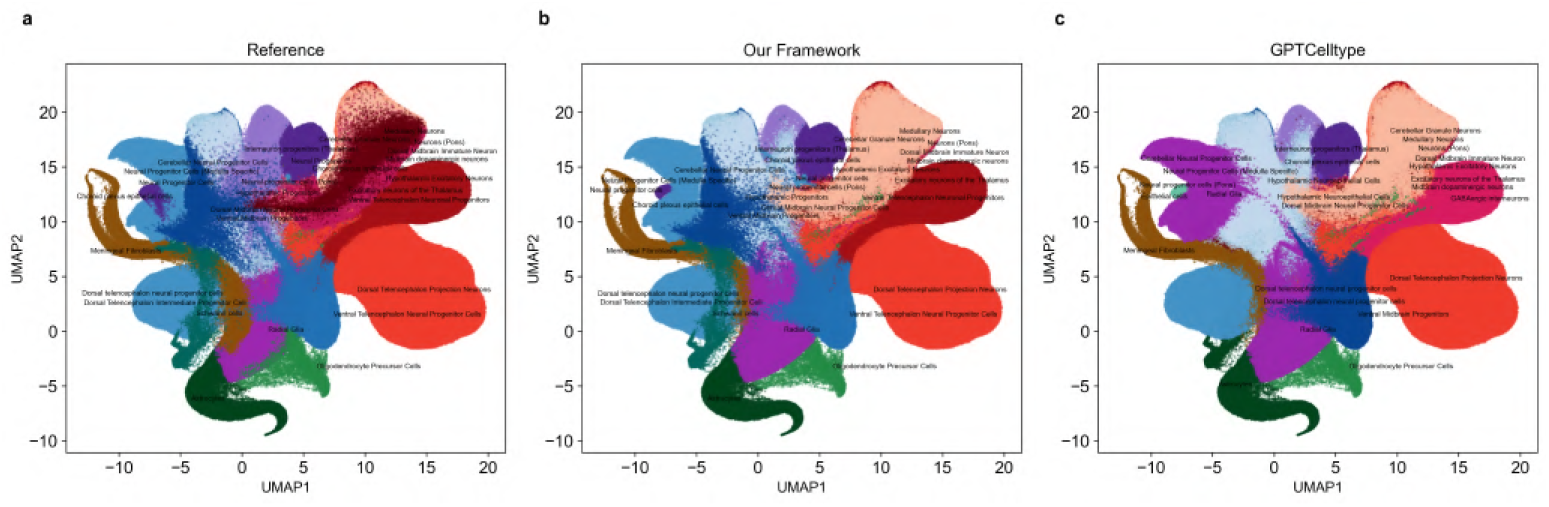
HNOCA dataset annotation results comparison. **a**, UMAP visualization of HNOCA reference annotations showing major neural cell type populations and developmental states. The reference annotations were generated through a multi-step process using the snapseed tool, which combines hierarchical cell type definitions, marker genes scoring, and reference mapping to human developmental brain atlases. The data was further refined through scPoli integration and label transfer from reference brain atlases using scVI and scANVI, particularly for non-telencephalic neurons and progenitor cells. **b**, UMAP visualization of our framework’s annotations demonstrating robust performance in identifying diverse neural cell types and developmental states, with high annotation accuracy when evaluated against reference annotations across this large-scale integrated atlas. **c**, UMAP visualization of GPTCelltype annotations showing lower accuracy compared to our framework, particularly in distinguishing between closely related neural progenitor states and specialized neuronal subtypes.

**Extended Data Figure 6:**
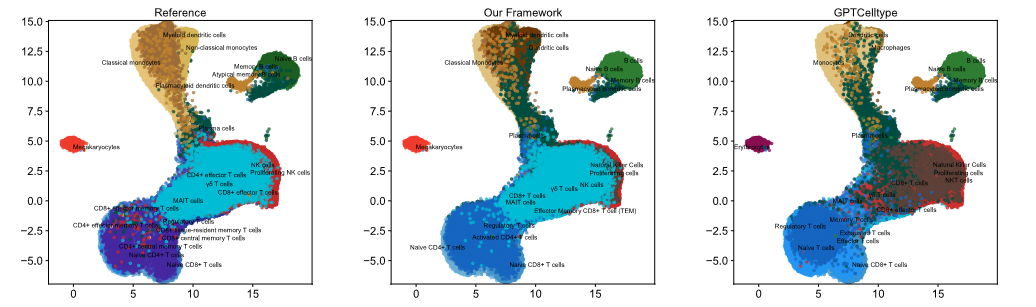
Annotation performance on the lifespan-wide human peripheral immune cell atlas. **a**, UMAP visualization of reference annotations showing the distribution of immune cell types across the human lifespan. The reference annotations were generated through a comprehensive analysis of immune cells from 220 healthy donors spanning 13 age groups from birth to over 90 years, combining transcriptomic profiles with expert annotation. **b**, UMAP visualization of our framework’s annotations demonstrating high annotation accuracy (76.6%) when evaluated against reference annotations, successfully capturing the complex developmental transitions and diverse immune cell populations.

**Extended Data Figure 7:**
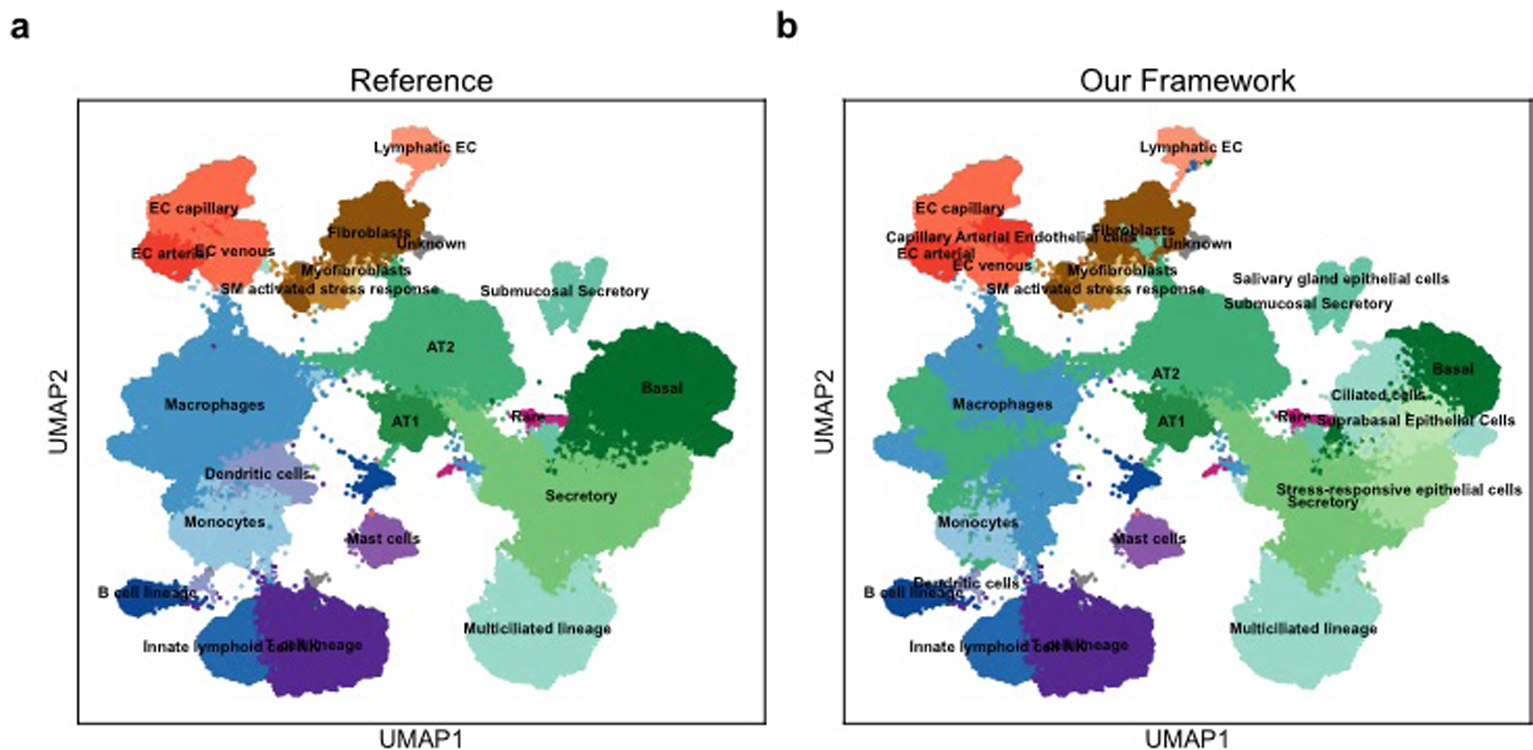
HLCA dataset annotation results comparison. **a**, UMAP visualization of HLCA reference annotations showing major cell type populations and their distribution. The HLCA reference annotations were generated through a hierarchical framework comprising 5 levels of granularity, from broad labels (Level 1: immune, epithelial, etc.) to fine-grained cell types (Level 5: e.g., naive CD4 T cells). The data integration and clustering were performed using scANVI, followed by Leiden clustering at different resolutions (Level 1: 0.01, Level 2: 0.2 with *k* = 30, Levels 3-5: 0.2 with *k* = 15*/*10). The final annotations were manually curated by six lung biology experts based on the clustering results, evidence from marker genes, and HLCA core clustering outcomes. **b**, UMAP visualization of our framework’s annotations demonstrating high annotation accuracy when evaluated against reference annotations. Following the same hierarchical clustering approach as the reference, our framework performed cell type annotation level by level. At each level, the LLMs were provided with global marker genes (genes specifically expressed in one cluster compared to all other clusters) and sister marker genes (genes differentially expressed between the target cluster and its sibling clusters within the same parent cluster), along with the parent cluster’s annotation from the previous level to guide the annotation process.

**Extended Data Figure 8:**
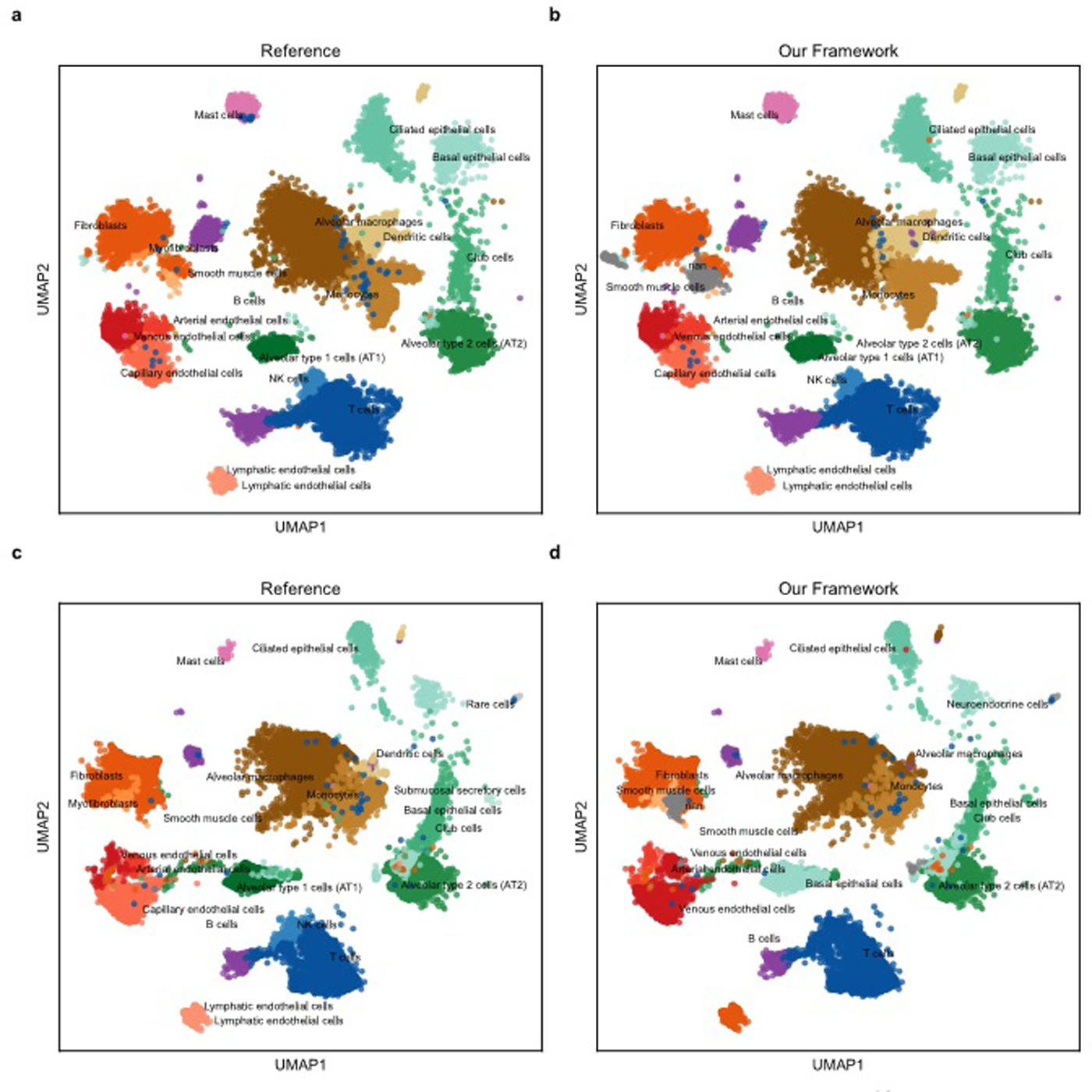
Our framework’s performance on alternative single-cell sequencing platforms. **a**, UMAP visualization of reference annotations from Drop-seq data,^12^ demonstrating the distribution of cell populations in this alternative sequencing platform. **b**, UMAP visualization of our framework’s annotations on Drop-seq data showing 95% annotation accuracy with reference annotations. **c**, UMAP visualization of reference annotations from snRNA-seq data,^13^ showing cell type distributions in the snRNA-seq platform. **d**, UMAP visualization of our framework’s annotations on snRNA-seq data demonstrating robust performance across different nuclear transcriptome profiles, validating our framework’s applicability to diverse scRNA-seq technologies.

**Extended Data Figure 9:**
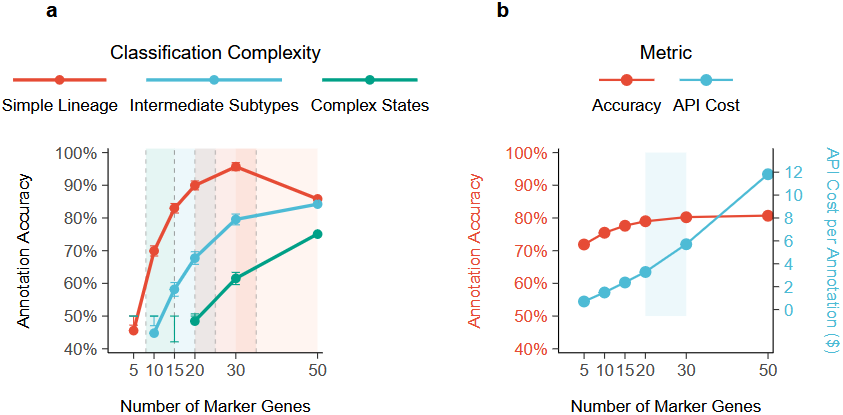
Optimization of marker genes selection for multi-LLM annotation framework. **a**, Quantitative assessment of annotation accuracy as a function of the number of marker genes across different classification complexities. Three distinct complexity levels are shown: simple lineage classification (red line, e.g., B/T cell distinction), intermediate subtypes (blue line, e.g., T cell subtypes such as CD4+/CD8+), and complex states (green line, e.g., naive/memory/effector states). Performance curves demonstrate distinct optimal ranges: simple lineage classification (8-15 marker genes), intermediate complexity (15-25 marker genes), and complex developmental states (20-35 marker genes). Each data point represents the annotation accuracy, with error bars showing standard deviation. The shaded region beyond 30 marker genes indicates potential diminishing returns and increased complexity. **b**, Dual analysis of framework efficiency versus marker genes count. The primary axis (red) shows annotation accuracy, while the secondary axis (blue) tracks API cost per annotation. The blue-shaded region (20-30 marker genes) highlights the optimal operational range that balances accuracy with resource utilization

**Extended Data Figure 10:**
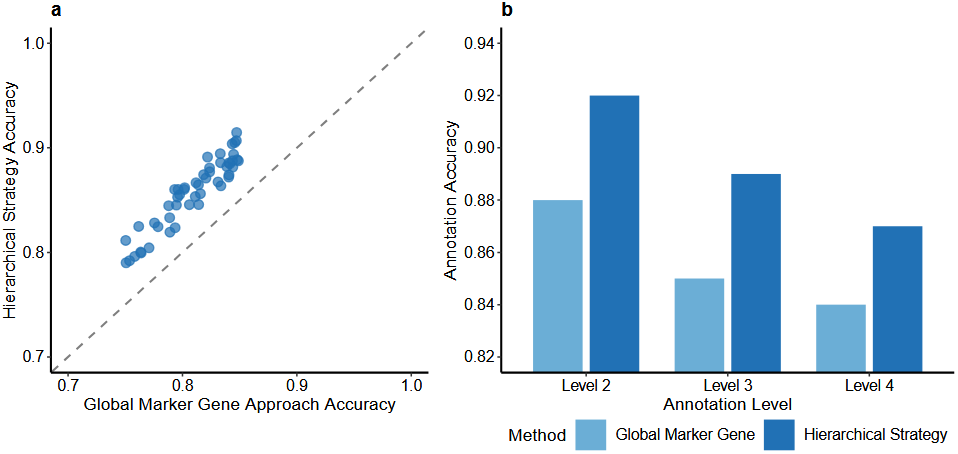
Impact of sister marker genes in hierarchical annotation strategy. **a**, Comparison of annotation results between two input strategies: using global marker genes (genes specifically expressed in one cluster compared to all other clusters) alone versus incorporating additional sister marker genes (genes differentially expressed between the target cluster and its sibling clusters within the same parent cluster). **b**, Quantitative analysis showing improvement in annotation accuracy when sister marker genes are added to the input alongside global marker genes. This improvement is particularly evident in distinguishing closely related cell types that share similar global expression patterns but differ in specific marker genes when compared to their sister clusters.

**Extended Data Figure 11:**
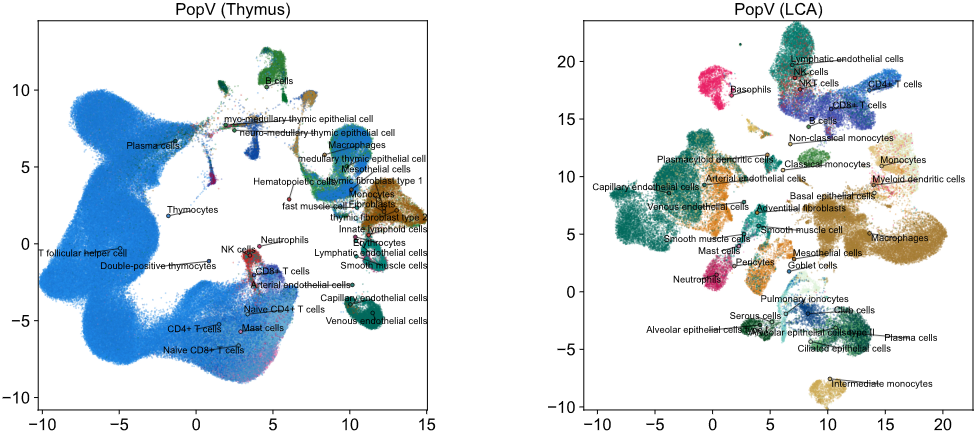
popV annotation performance on benchmark datasets. **a**,MAP visualization of popV annotations on the developmental human thymus dataset, showing limitations in accurately distinguishing between closely related T cell developmental stages and specialized thymic cell populations. **b**, UMAP visualization of popV annotations on the Lung Cell Atlas (LCA) dataset, demonstrating challenges in correctly identifying specialized lung cell subtypes and rare cell populations.

## 6 Supplementary Information

Supplementary information is available for this paper at [URL will be provided by the journal].

Supplementary information includes:

- Supplementary Table 1: Complete collection of prompt templates used in our multi LLM consensus framework, organized by function (initial annotation, deliberation, consensus checking) with full prompt text and explanations of key components
- Supplementary Table 2: Comprehensive comparison of mLLMCelltype with other representative scRNA-seq cell type annotation methods, highlighting methodological differences, input requirements, reference dependencies, knowledge integration approaches, uncertainty quantification capabilities, scalability characteristics, novel cell type detection potential, and key advantages and limitations of each method

## 7 Acknowledgements

We acknowledge the publicly available datasets used in our benchmarking, including those from Tabula Sapiens, Human Cell Landscape, Mouse Cell Atlas, Genotype Tissue Expression (GTEx), Human BioMolecular Atlas Program (HuBMAP), Human Lung Cell Atlas (HLCA), Human Neural Organoid Cell Atlas (HNOCA), the lifespan wide human peripheral immune cell atlas, and specialized datasets including B cell lymphoma (BCL) and cancer datasets from colon and lung. We also thank the developers of the open-source tools utilized in this work.

## 8 Declarations

### 8.1 Ethics approval and consent to participate

This study relies solely on publicly available datasets, for which ethical approvals and participant consents were obtained as reported in the original studies. No additional ethical approval was required for our analysis.

### 8.2 Funding

The work is supported by the National Institute of Health R01GM144351 (Chen & Zhang), National Science Foundation DMS1830392, DMS2113359, DMS1811747 (Zhang), National Science Foundation DMS2113360, and Mayo Clinic Center for Individualized Medicine (Chen).

### 8.3 Conflict of interest/Competing interests

The authors declare no competing interests.

### 8.4 Consent for publication

Not applicable.

### 8.5 Materials availability

No physical materials were used in this study. Software and data availability are addressed in their respective sections.

### 8.6 Author contribution

C.Y., X.Z., and J.C. jointly conceptualized the study. C.Y. developed the methodology, implemented the computational framework, conducted formal analysis, and performed data curation, visualization, and validation. J.C. contributed to the experimental design. X.Z. and J.C. jointly supervised the research, secured funding, provided resources, served as corresponding authors, and administered the project, contributing equally in these senior roles. C.Y., X.Z., and J.C. all participated in manuscript writing and revision.

## Notes

https://cells.ucsc.edu/?ds=tabula-sapiens

https://figshare.com/articles/dataset/HCL_DGE_Data/7235471

https://figshare.com/s/865e694ad06d5857db4b

https://cellxgene.cziscience.com/e/9f222629-9e39-47d0-b83f-e08d610c7479.cxg/

https://cellxgene.cziscience.com/collections/5d445965-6f1a-4b68-ba3a-b8f765155d3a

https://cellxgene.cziscience.com/collections/de13e3e2-23b6-40ed-a413-e9e12d7d3910

https://cellxgene.cziscience.com/collections/de379e5f-52d0-498c-9801-0f850823c847

https://www.synapse.org/Synapse:syn61609846

https://azimuth.hubmapconsortium.org/

https://gtexportal.org/home/datasets

### Competing Interest Statement

The authors have declared no competing interest.

