## Supplementary Table 2 for "Large Language Model Consensus Substantially Improves the Cell Type Annotation Accuracy for scRNA-seq Data"

Table 1: **Supplementary Table 2 — Comprehensive comparison of mLLMCelltype with other representative scRNA-seq cell type annotation methods.** This table provides a detailed feature-by-feature comparison highlighting the methodological differences, input requirements, reference dependencies, knowledge integration approaches, uncertainty quantification capabilities, scalability characteristics, novel cell type detection potential, and key advantages and limitations of each method.

| Feature | mLLMCelltype (Proposed) | GPTCelltype | popV | SingleR | scCATCH |
|---|---|---|---|---|---|
| **Methodology** | Multi-LLM Consensus Deliberation | Single LLM (GPT-4 based) | Ensemble Learning (Multiple algorithms) | Supervised Correlation (Cell-level vs Reference) | Unsupervised + Marker DB (Cluster-level vs DB) |
| **Input Required** | Cluster marker genes, Tissue context | Cluster marker genes, Tissue context | Cell expression matrix, Tissue context (optional), Pre-trained model / Ref. | Cell expression matrix, Annotated reference expression dataset | Clustered data, Tissue context (for DB) |
| **Reference Dependent?** | No (uses LLM knowledge) | No (uses LLM knowledge) | Yes (for pre-training or retraining) | Yes (Requires annotated expression reference) | No (Requires marker database, not expression ref.) |
| **Knowledge Integration** | Implicit (LLM training) + Explicit (Deliberation) | Implicit (LLM training) | Data-driven ensemble (Implicit in selection) | None explicit | Explicit (Marker DB) |
| **Uncertainty Quant.** | Yes (Consensus Prop., Shannon Entropy) | Limited (Reasoning only) | Yes (Ensemble scores) | Yes (Correlation scores) | Limited (Matching scores) |
| **Scalability** | High (Cluster-level, API cost scales) | High (Cluster-level, API cost scales) | Moderate-High (GPU often needed for retraining) | Moderate (Cell-by-cell comparison) | High (Cluster-level) |
| **Novel Type Detection** | Potential (via LLM inference, deliberation) | Potential (via LLM inference) | Limited (by ensemble components/refere | No (Assigns best match from reference) | Limited (Requires clustering + DB match) |
| **Key Advantage** | High accuracy, Robustness, Uncertainty Quant., Reasoning Transparency | High automation, Leverages LLM knowledge | High accuracy (w/ good ref.), Ensemble robustness | Simple, Widely used | No annotated expr. ref. needed, Uses known markers |
| **Key Limitation** | API costs, Potential collective LLM bias | Single model bias, Lacks formal uncertainty | Reference-dependent, Computationally intensive | Highly reference-dependent, Cannot find novel types | Marker DB quality/coverage, Clustering dependent |
| **Automation Level** | High (Flags uncertain) | High | High (Post-training) | High | High (Post-clustering) |
